# Thermal environment and ecological interactions modulate the importance of evolution in response to warming

**DOI:** 10.1101/2023.09.28.559813

**Authors:** Cara A. Faillace, Soraya Álvarez-Codesal, Alexandre Garreau, Elvire Bestion, José M. Montoya

## Abstract

Understanding the interaction between evolutionary history, the current abiotic environment, and biotic interactions is critical for a more nuanced understanding of the response of communities to anthropogenic stressors. We leveraged a long term experiment manipulating temperature in mesocosms containing communities of phytoplankton and zooplankton to examine how evolution in response to long-term community warming affects consumer-resource dynamics at different temperatures. We showed that the evolution in response to warming depends on both the current thermal environment, as well as the presence of interactions between consumers and resources. We also demonstrated that evolution influences the outcomes of current ecological dynamics. For each consumer–resource pair, the effects of evolution were temperature-dependent, but both the effects and the temperature dependence itself additionally depended upon the identity of evolving species in each pair. Evolution resulted on a win-win situation for the first resource species: across all temperatures, this resource was more fit and the consumer was less successful, with fitness gains peaking at intermediate temperatures. For this resource species our results supported the “hotter is better” hypothesis, especially at moderate or intermediate temperatures, while “hotter is worse” for the consumer. In the second species pair, patterns were more complex. Warm-origin populations of both the second resource and the consumer generally failed to show improved fitness. Overall, our results show that evolution altered resource and consumer fitness, but these effects were dependent on the current combination of abiotic and biotic conditions.

## Introduction

Climate change has emerged as a critical challenge to species and ecological communities (Parmesan 2006, Radchuk et al. 2019). Multiple mechanisms by which species and communities respond to increasing temperatures (e.g., range shifts, plastic changes to behavior, phenology, etc.) interact to determine the degree to which species and communities are affected by climate change (O’Connor et al. 2012, Lurgi et al. 2012, Gaitán-Espitia and Hobday 2021). Because temperature represents a strong selective agent for populations, evolution likely constitutes one important mechanism by which species respond to warming, especially for dispersal-limited populations or species with rapid generation times (O’Donnell et al. 2018). Evolution in response to warming has been experimentally documented in a variety of organisms, including, for instance, green algae (Padfield et al. 2016, Schaum et al. 2017), arthropods (Gilchrist and Huey 1999, Geerts et al. 2015, Hangartner and Hoffmann 2016), and larger organisms, such as lizards (Logan et al. 2014). While less frequently studied, evolution in natural populations in response to warming has been observed as well (Higgins et al. 2014, Geerts et al. 2015, Johansson et al. 2016, Johansson and Laurila 2017, Diamond et al. 2017). Because evolution in response to abiotic factors, like climate warming, is frequently explored in single populations, the implications of evolution in communities responding to warming remain understudied. Nonetheless, given that many of the species most likely to evolve in response to warming—such as plankton with short generation times and larger population sizes—are also critical for ecosystem functioning and services (*e.g.,* water quality, carbon sequestration)(O’Donnell et al. 2018), we can expect that evolutionary responses to warming will have important consequences for both ecological and human systems (Rudman et al. 2017).

Multiple hypotheses exist for how evolution in response to warming may alter the populations’ thermal response curves of a number of fitness attributes [e.g., “hotter is better”, “biochemical adaptation”, and “partial compensation” (Angilletta et al. 2010, Liu et al. 2022)]. Evidence from natural populations, however, demonstrates that the thermal performance of populations can depart from theoretical expectations (DeLong et al. 2018). In some cases responses match anticipated shifts in thermal performance with adaptation under warmer temperatures (Knies et al. 2006, Diamond et al. 2017). In other cases they exhibit unexpected results, which can also depend upon the life history parameter measured (Kingsolver and Huey 2008, Johansson et al. 2016). Such unforeseen results may, in part, be explained by life history tradeoffs (Latimer et al. 2011), such as those that can emerge between coping with acute thermal stress and chronic exposure (Rezende et al. 2014). For instance, evolution may favor increased survival under chronic warm temperatures at the expense of critical thermal maximum temperature during acute thermal stress (Johansson and Laurila 2017). Thus, both the type of thermal stress experienced by populations (i.e., increasing mean temperature vs. frequent acute fluctuations), as well as the current thermal environment (e.g., temperature and duration of thermal experiments) may be important for interpreting empirical patterns and understanding the implications of evolution that occurs in response to warming.

The presence of additional interacting species introduces additional factors influencing evolutionary outcomes. The effects of evolution can depend on the current ecological conditions of populations experiencing warming. When species are embedded in complex communities, the opportunities for additional costs, constraints, or benefits of evolution can emerge. For instance, there can be trait tradeoffs or novel selective pressures present that arise out of additional ecological (or evolutionary) changes that occur as communities respond to warming. Such trait tradeoffs can drive differences in evolutionary responses compared to evolution in single populations (Van Doorslaer et al. 2009a, 2010). Selection that occurs within an ecological community is therefore likely to qualitatively change predictions about thermal adaptation derived from single populations (Angilletta Jr. et al. 2006), such that individual species’ responses to temperature can be insufficient to explain changes to community composition (Tabi et al. 2020).

In consequence, we expect that when shifts in thermal performance occur in traits that affect how species interact, such as resource acquisition or metabolism, this could affect overall fitness of populations that interact and respond to warming. Specifically, 1) different life history traits within a species can have different thermal dependencies (Dell et al. 2011, Huey and Kingsolver 2011), 2) consumer and resource species have documented differences in these thermal dependencies (O’Connor et al. 2009, Álvarez-Codesal et al. 2023), and 3) evolutionary potential can differ across trophic levels (Terhorst et al. 2010, Hague and Routman 2016). Therefore, it follows that evolution in response to warming has the potential to alter consumer-resource dynamics, while also being reciprocally influenced by them (Faillace et al. 2021), and that the effects of evolution may additionally depend upon current thermal environment. For example, previous research using a space-for-time substitution demonstrated that locally adapted populations of consumer and resource species from different latitudes had contrasting eco-evolutionary dynamics at two temperatures (De Block et al. 2013). More generally, a variety of altered dynamics are possible. One possibility that emerges is cryptic eco-evolutionary dynamics where evolution in both species yields a “moving target” that mimics purely ecological dynamics. In this case, evolution may alter the thermal performance of the interacting species in similar ways, such that the outcomes of the interaction remain apparently unchanged, but are actually being maintained by the evolutionary response. A second possibility is that the effects of the interaction on species’ fitness may depend upon the current thermal environment. This might occur if, for example, their thermal tolerances evolve to different degrees or if only one interacting species evolves. Additionally, although responses are sometimes examined at multiple temperatures, most studies are typically of short duration and do not consider the effects of evolution at longer timescales to quantify changes to fitness.

To examine both the influence of ecological interactions on evolutionary responses to warming as well as understand the long-term effects of evolution on the ecology of interacting species, we used a long-term warming mesocosm experiment in plankton communities. We examined the consequences of evolution in response to long-term warming in communities for consumer-resource interactions. We used populations of two resource species (i.e., freshwater green algae, *Chlamydomonas reinhardtii* and *Desmodesmus* sp.) and a consumer species (i.e., *Daphnia pulex*) differing in thermal origin. We hypothesized that the effects of evolution on species’ fitness would be dependent upon current thermal environment and the presence of species interactions. Specifically, we asked, 1) whether biotic interactions affect the importance of evolutionary history across a range of temperatures, and 2) whether temperature affects the importance of evolutionary history.

## Methods

### Source and maintenance of organisms prior to experimentation

Organisms originated from a long-running mesocosm experiment in Dorset, England. This mesocosm experiment has been the subject of multiple previous publications (e.g., Yvon-Durocher et al. 2015, Schaum et al. 2017, Álvarez-Codesal et al. 2023). Briefly, at the time of sampling in 2019, open-air experimental ponds of 1 m^3^ water of mixed freshwater phytoplankton and zooplankton had been continuously maintained since 2005 (i.e., approximately 14 years). Half of the mesocosms were kept at ambient temperature conditions, while half experienced a consistent temperature increase of +4 °C above ambient temperatures (n = 8 for both at the time of our sampling). Temperature in all ponds fluctuated seasonally. Mesocosm communities initially contained the same mixture of phytoplankton and zooplankton, but previous work has demonstrated significant changes to community structure in the warmed mesocosms, as well as evolution in thermal tolerance in warmed populations of at least one phytoplankton species, *Chlamydomonas reinhardtii* (Yvon-Durocher et al. 2015, Schaum et al. 2017)(Figure 1).

**Figure 1.**
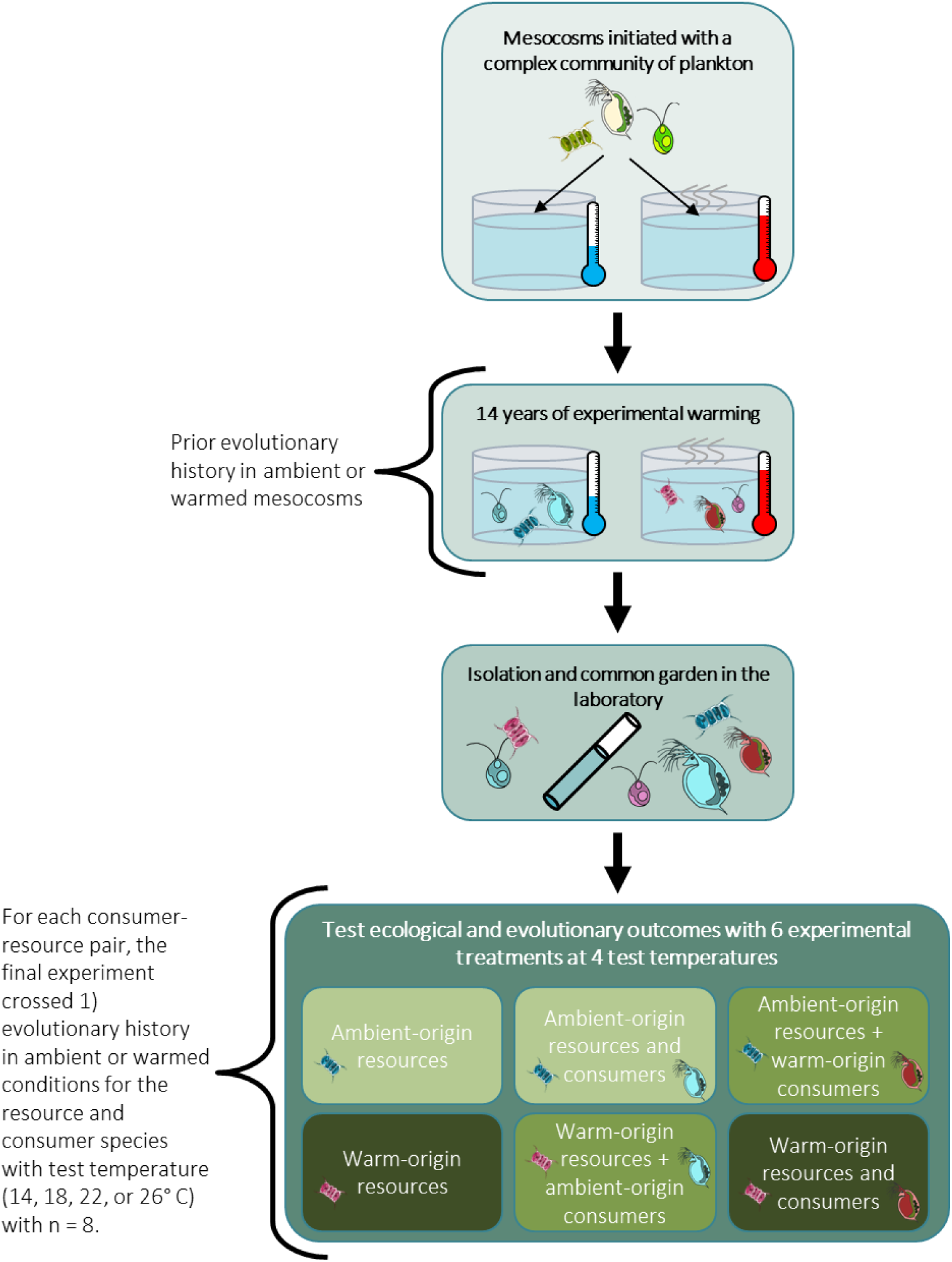
Experimental design in which populations from two resource species and one consumer species were first collected from a long-running warming experiment using communities of plankton in outdoor mesocosms. After isolation and a period of laboratory common garden, for each consumer– resource pair, subsequent experiments examined the interaction between evolutionary history of resources and consumers and current thermal environment on population abundance.

We sampled two phytoplankton resource species, the algae *C. reinhardtii* and *Desmodesmus* sp. from each treatment in January 2019, as well as a consumer species, *Daphnia pulex* in October 2019 (for further details see Álvarez-Codesal et al. 2023). The discrepancy in dates occurred because we were unsuccessful in isolating consumers on the first attempt. Sorting flow cytometry isolated single algal cells and PacBio Sequel 2 long-read sequencing using full-length 18S fusion primers (forward: NSF4/18 = CTGGTTGATYCTGCCAGT; reverse: EukR = TGATCCTTCTGCAGGTTCACCTAC) confirmed species identity and genotypic differences between ambient-origin and warm-origin populations of the algae (Comeau et al. 2017). We identified one unique clone for each of the two species, *C. reinhardtii* and *Desmodesmus* sp. The identity of the consumer populations originating from each of 5 individuals isolated from the experiment was confirmed morphologically by a taxonomist (however, genotypic differences were not verified by sequencing for the *Daphnia*). All species reproduced asexually, enabling us to maintain clonal populations of each genotype (algae) or putative clone (*Daphnia*) in semi-continuous laboratory batch cultures until the start of the experiments. To avoid confounding effects of plasticity or maternal effects, we maintained populations of these species in common garden conditions—20 °C with 12:12 light-dark cycle—for over six months (many generations) prior to subsequent experimentation. All organisms grew in autoclave-sterilized COMBO growth medium (Kilham et al. 1998). *Daphnia* grew in modified low-phosphorus COMBO (Kilham et al. 1998) that reduces bacterial load. Aeration maintained culture health and growth, and we aimed to maintain algal populations in the exponential growth phase leading up to the experiment. Additionally, *D. pulex* grew on a laboratory culture of *Ankistrodesmus* sp. (sourced from Sciento UK) to ensure that daphnids were not acclimated to the algal species used in the experiments. *Daphnia* cultures received 50 mL of dense *Ankistrodesmus* once per week as a food source. Periodic filtering of adult and juvenile daphnids into fresh medium removed metabolic waste products and resting eggs.

### Experimental design

Our experimental design crossed long-term evolutionary history—either to “ambient” or “warm” temperature conditions— of each consumer–resource pair with 4 growth temperatures (Figure 1). In this way, we sought to examine how the effects of evolution in response to warming in a consumer–resource pair depend upon the current thermal environment. The experiment occurred in 4 incubators that varied in temperature: 14, 18, 22, and 26 °C. These temperatures encompass much of the observed growing season activity range for the consumer from this system. The presumed approximate thermal optimum of reproduction of the ambient-origin population of 20 °C served as the mid-point across the range (Craddock 1976). Non-consumption controls additionally allowed us to verify the importance of algal evolutionary history in the absence of consumers. We conducted the experiment for each consumer–resource pair in two temporal blocks, across which we swapped incubator temperatures (for tractability and to avoid pseudoreplication at the incubator level). Thus, we had 6 warming evolutionary history–consumption treatments crossed with 4 growth temperatures (Figure 1) in each temporal block for each consumer–resource pair (n = 4 per treatment, yielding a total of 96 microcosms within temporal block). At the start of each experimental block, we estimated abundance of the ambient-origin and warm-origin resources using a hemocytometer to count cells under a compound microscope (NIKON Eclipse Ni). We then standardized approximate starting abundances of the two populations, however starting densities thus differed somewhat between blocks within a resource species and between resource species. We randomized microcosm position within each incubator. The experiment ran a total of 4 times (n = 8 for treatment across 2 temporal blocks, yielding N = 192 per consumer–resource pair and N = 384 across both pairs). The experiment ran for 3 weeks, or approximately 42–100 generations of algae and 2–4 generations of *Daphnia*.

To each sterile microcosm we added 100 mL of sterile COMBO medium and 100 mL of density-standardized resource from either the ambient-origin or warm-origin population (of either *C. reinhardtii* or *Desmodesmus* sp.). To treatments containing *Daphnia*, we then added 5 daphnids either of ambient-origin or warm-origin to each microcosm, one from each of 5 isoclonal lineages of ambient-origin or warm-origin populations of *Daphnia*, such that the evolutionary origin of the resource was crossed with that of the consumer. It was infeasible to grow out the number of juveniles that would be needed to ensure precisely aged populations from the 5 isoclonal *Daphnia* lineages from both the ambient- and warm-origin populations. Therefore, we chose from among largest adult daphnids to ensure that we included individuals of the largest instar for each population. This provided a snapshot of the *Daphnia* community at the point at which we initiated the experiment, but the adults may have differed in age, contributing to differences among temporal blocks. The *Daphnia* were previously starved for 24-hours, filtered, and rinsed in sterile water to remove residual *Ankistrodesmus* before addition to the experiment. Thus our evolutionary history–consumption treatments repeated at 4 temperatures were: 1) ambient-origin resource alone, 2) warm-origin resource alone, 3) both ambient-origin resource and consumer, 4) ambient-origin resource + warm-origin consumer, 5) warm-origin resource + ambient-origin consumer, and 6) both warm-origin resource and consumer.

### Estimation of equilibrium densities and interaction strength

Responses included equilibrium density (per ml) of the resource species, as well as a census of adult and larval *Daphnia* present in each microcosm. After 3 weeks, we first filtered and preserved all *Daphnia* adults and juveniles from a microcosm in 95% ethanol for later enumeration under a dissecting scope in two size categories, adults and juveniles. Safranin (Sigma-Aldrich) stained the *Daphnia* for simpler visualization of the preserved samples during counting under a stereo microscope (Leica S9i), which ensured that molts were not counted erroneously. We next cryopreserved two 150 μL samples of well-mixed algae suspension from each microcosm using 15 μL of a 1% sorbitol solution in 96-well plates at –80 °C. Flow cytometry (BD FACSCanto II high-throughput sampler) then estimated cell densities (for more details, see Álvarez-Codesal et al. 2023). For *Desmodesmus* sp., because flow cytometry measures particles, we report our results as coenobia (i.e., small colonial cellular aggregates) rather than converting from aggregates to cells. Coenobia are the units with which consumers interact, making them an appropriate unit for this species.

We used interaction strength to quantify the effect of consumers on resource abundance. To estimate the population-level interaction strength (IS), we calculated an inverse form of the log-response ratio (also called the Dynamic Index) using Equation 1 (Berlow et al. 2004, Schaum et al. 2017):

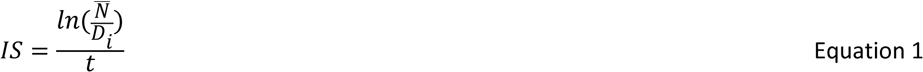

where 𝑁̅ is the final mean algal abundance of the no consumption (algae-alone) controls for each population, ambient-origin or warm-origin, for each resource species, *C. reinhardtii* or *Desmodesmus* sp., 𝐷_𝑖_ is the final algal abundance of replicate 𝑖 from a given population in the presence of consumers, and 𝑡𝑡 is the length of the experiment (equal to 21 days for our experiment). Larger values of IS indicate stronger effects of consumers on resource abundance. We chose to focus on population-level interaction IS rather than the *per capita* IS because we measured the interaction over a period of 21 days (i.e., it was not a short-term measure) during which the consumer population size was not constant. Thus, we might have some consumer individuals dying and others hatching and emerging during the experiment, so that populations are not at equilibrium, making the population-level log-response ratio an appropriate choice for our design (Berlow et al. 2004).

### Analyses

We log_10_-transformed abundances to better meet assumptions of normality in the subsequent analyses. In all models, a block effect and all block interactions accounted for differences related to temporal blocks. In all analyses, we centered the temporal blocking variable to ensure interpretability (Schielzeth 2010). We conducted analyses to examine how current thermal environment and biotic interactions determined the importance of past evolutionary history to warming in governing fitness (as assessed by abundance across multiple generations) of resources and consumers. A single MANOVA including evolutionary history treatments with and without consumers present would obfuscate differences that might occur in the interaction between evolutionary history and temperature when the algae grew alone or with consumers (in part through spurious significance related to the presence or absence of *Daphnia* as a response in the two types of treatments). Therefore, to examine how current biotic interactions impact the importance of evolutionary history, we conducted separate analyses on the log_10_-transformed abundance per ml of algae when consumers were present or absent. In all models, the interaction between the main effects (evolutionary history and temperature) assessed whether the effects or importance of evolutionary history depend upon current thermal environment. Thus, we first conducted an analysis of variance (ANOVA) for each algal species grown alone (because they were tested independently). These analyses tested whether algal evolutionary history interacted significantly (α = 0.05) with temperature when consumers were absent. The model included the evolutionary history (2 levels—ambient-origin or warm-origin resources) and temperature (4 levels— 14, 18, 22, and 26 °C test temperature) and the interaction, in addition to the block effect and block interactions, with final algal density as the response.

We next conducted a separate multivariate analysis of variance (MANOVA) for each consumer– resource pair. Responses included the log_10_-transformed abundance per ml of the resource, interaction strength, and the log_10_-transformed abundances per microcosm of adult and juvenile *Daphnia*. Main effects of the model included evolutionary history treatment (4 levels—ambient-origin resource and consumers, ambient-origin resource + warm-origin consumers, warm-origin resource + ambient-origin consumers, and warm-origin resource and consumers), temperature (4 levels—14, 18, 22, and 26 °C test temperature), and the interaction between temperature and evolutionary history. Univariate tests assessed significance at α = 0.05 for individual responses. Planned contrasts further disentangled the drivers underlying the evolutionary history treatment and determined specifically if 1) resource evolution, 2) consumer evolution, or 3) the interaction between evolutionary history of the resource and consumer (i.e., coevolution) affected responses. All analyses were conducted in SAS 9.4 (SAS Institute 2011).

## Results

Evolutionary history in response to warming in complex communities, current temperature, and their interaction had highly significant effects for both consumer–resource pairs, but the relative importance of resource, consumer, or coevolution differed between the pairs (see Tables 1 and 2 for specific results from algae-alone ANOVAs and consumer–resource MANOVAs). We describe specific effects below using results from ANOVAs and planned contrasts for each response variable.

**Table 1.** Results from univariate ANOVAs (n = 8) examining the effects of resource evolution when growing alone for *Chlamydomonas reinhardtii* and *Desmodesmus* sp. The models included an evolution treatment and temperature as two main effects, as well as a temporal blocking variable and all interactions. Here, the evolution treatment included just the evolutionary history of the resource species, either ambient-origin or warm-origin. The interaction between evolution treatment and temperature evaluated any temperature-dependence of the effects of evolution. Significant results are in bold.

| *C. reinhardtii* |  |  |  |
| --- | --- | --- | --- |
|  | d.f. | F-value | p-value |
| **Evolution** | 1 | **158.47** | **< 0.0001** |
| **Temperature** | 3 | **12.56** | **< 0.0001** |
| **Temporal block** | 1 | **54.12** | **< 0.0001** |
| Evolution–temperature | 3 | 0.67 | 0.5776 |
| **Temperature–block** | 3 | **15.43** | **< 0.0001** |
| **Evolution– block** | 1 | **127.63** | **< 0.0001** |
| **Evolution–temperature–block** | 3 | **5.96** | **0.0015** |
| *Desmodesmus* sp. |  |  |  |
|  | d.f. | F-value | p-value |
| **Evolution** | 1 | **7.02** | **0.0109** |
| **Temperature** | 3 | **47.41** | **< 0.0001** |
| Temporal block | 1 | 3.94 | 0.0529 |
| **Evolution– temperature** | 3 | **5.86** | **0.0017** |
| **Temperature–block** | 3 | **5.19** | **0.0035** |
| **Evolution–block** | 1 | **42.75** | **< 0.0001** |
| **Evolution–temperature–block** | 3 | **3.10** | **0.0353** |

**Table 2.** MANOVA (n = 8) results examining the effects of evolution of both the consumer and the resource in complex communities in response to warming for two consumer–resource pairs, *C. reinhardtii* and *D. pulex*, and *Desmodesmus* sp. and *D. pulex*. The models included an evolution treatment and temperature as two main effects, as well as a temporal blocking variable and all interactions. Here, the evolution treatment included the four possible combinations crossing the ambient- or warm-origin populations of both resources and consumers, but these analyses did not include algae-alone controls. The interaction between the evolution treatment and temperature evaluated any temperature-dependence of the effects of evolution. Orthogonal contrasts evaluated the effects of 1) resource evolution, 2) consumer evolution, and 3) coevolution between the resource and the consumer. Reported are Wilks’ Lambda F-values and associated p-values. Significant results are in bold.

| *C. reinhardtii*–*D. pulex* |  |  |  |
| --- | --- | --- | --- |
|  | d.f. | F-value | p-value |
| **Evolution** | 12, 243.7 | **70495.6** | **< 0.0001** |
| **Temperature** | 12, 243.7 | **45724.4** | **< 0.0001** |
| **Temporal block** | 4, 92 | **1.82E10** | **< 0.0001** |
| **Evolution–temperature** | 36, 346.5 | **1163.05** | **< 0.0001** |
| **Temperature–block** | 12, 243.7 | **51809.9** | **< 0.0001** |
| **Evolution– block** | 12, 243.7 | **63495.4** | **< 0.0001** |
| **Evolution–temperature–block** | 36, 346.5 | **1863.05** | **< 0.0001** |
| **Resource evolution contrast** | 4, 92 | **4.93E10** | **< 0.0001** |
| Consumer evolution contrast | 3, 93 | 1.72 | 0.1683 |
| Coevolution contrast | 3, 93 | 1.05 | 0.3729 |
| *Desmodesmus* sp.–*D. pulex* |  |  |  |
|  | d.f. | F-value | p-value |
| **Evolution** | 12, 243.7 | **578.58** | **< 0.0001** |
| **Temperature** | 12, 243.7 | **563.64** | **< 0.0001** |
| **Temporal block** | 4, 92 | **2818757** | **< 0.0001** |
| **Evolution–temperature** | 36, 346.5 | **56.96** | **< 0.0001** |
| **Temperature–block** | 12, 243.7 | **363.01** | **< 0.0001** |
| **Evolution– block** | 12, 243.7 | **633.42** | **< 0.0001** |
| **Evolution–temperature–block** | 36, 346.5 | **61.15** | **< 0.0001** |
| **Resource evolution contrast** | 4, 92 | **55119.7** | **< 0.0001** |
| **Consumer evolution contrast** | 3, 93 | **52771.4** | **< 0.0001** |
| **Coevolution contrast** | 3, 93 | **55945.4** | **< 0.0001** |

### Current thermal environment influences the importance of prior resource evolution in response to warming

Resource evolution in warmed communities affected both algal species when grown alone, but the effects of evolutionary history were opposite in direction and differed in strength (Table 1). Warm-origin *Chlamydomonas reinhardtii* had higher abundance than the ambient-origin genotype at all temperatures with no significant interaction between resource evolutionary history and current thermal environment (effect of evolutionary history in algae-alone ANOVAs: F_1_ = 158.47, p < 0.0001; interaction between evolutionary history and temperature: NS), an effect that appeared to be driven by differences across resource evolutionary history in temporal block 2 (Figure 2A, Figures S1 and S2). For *Desmodesmus* sp., the effect of evolutionary history depended upon temperature, with higher abundance of the warm-origin genotype at 14 °C, but lower abundance than the ambient-adapted genotype at the remaining three temperatures (effect of evolutionary history in algae-alone ANOVAs: F_1_ = 7.02, p = 0.0109; interaction between evolutionary history and temperature: F_3_ = 5.86, p = 0.0017) (Figure 2B, Figure S1). Here, higher abundance of the warm genotype at 14 °C was observed in temporal block 1, while the reduced abundance of warm genotype was more generally observed in temporal block 2 (Figure S2).

**Figure 2A–D.**
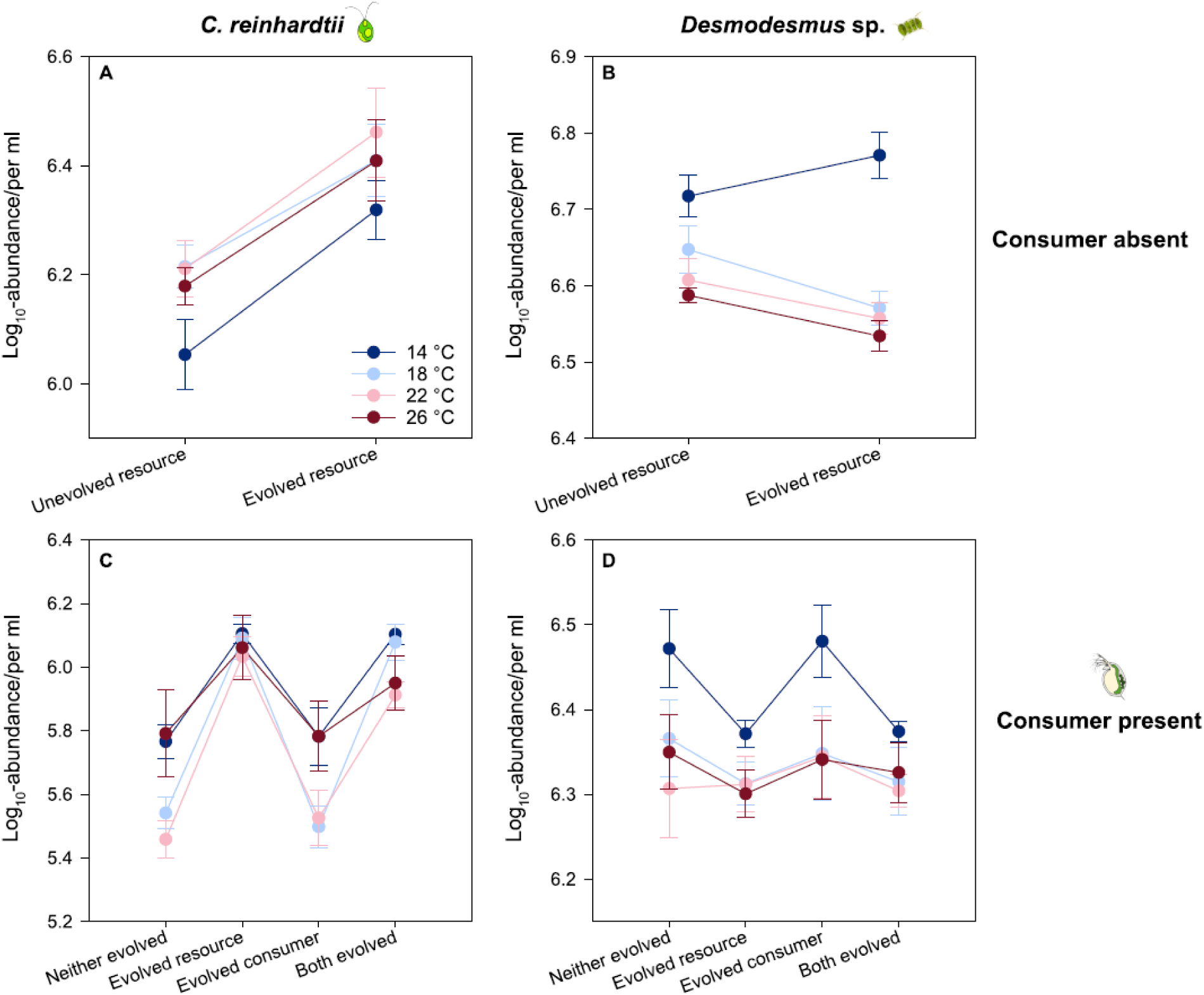
The effects of evolution in warmed communities on the log_10_-abundance of *C. reinhardtii* and *Desmodesmus* sp. depend upon current thermal environment and whether consumers are absent (A, B) or present (C, D), with related effects to the consumer–resource interaction strength (E, F). Evolutionary history treatments (where “evolved” refers to evolution in the warmed communities) are shown on the x-axis. Circles are the means (n = 8) of each treatment with the error bars showing ± the SEM. Colors correspond to temperature (dark blue = 14, light blue = 18, light pink = 22, and dark pink = 26 °C).

### Interspecific interactions modulate how current thermal environment interacts with prior evolution in response to warming Effects of interspecific interactions on resource abundance

The presence of the consumer *Daphnia pulex* altered the effects of resource evolutionary history for both resources, but in opposite directions (Tables 2–4). For *C. reinhardtii*, where no significant interaction between resource evolutionary history and temperature occurred when grown alone, the presence of the consumer resulted in a significant evolutionary history by temperature interaction driven by resource evolution (consumer–resource ANOVA resource evolution contrast: F_1_ = 318.47, p < 0.0001). Specifically, when consumers were present, the importance of resource evolutionary history was higher at intermediate temperatures (18 and 22 °C) than at the two temperature extremes (14 and 26 °C) (Figure 2C). In other words, temperature did not affect the direction of the response to evolutionary history. It did, however, increase the magnitude of the effect size at intermediate compared to extreme temperatures (effect of evolutionary history in consumer– resource ANOVA: F_3_ = 107.54, p < 0.0001; interaction between evolutionary history and temperature: F_9_ = 4.93, p < 0.0001). The temperature dependence of evolutionary history was stronger in the first temporal block (Figure S2). On the other hand, for *Desmodesmus* sp., the presence of consumers rendered the interaction between resource evolutionary history and temperature non-significant (effect of evolutionary history in consumer–resource ANOVA: F_3_ = 3.54, p = 0.0175; interaction between evolutionary history and temperature: NS) (Figure 2D). Here, when consumers were present, resource evolution decreased algal abundance, reversing the pattern seen at 14 °C in the absence of consumers (consumer–resource ANOVA resource evolution contrast: F_1_ = 10.57, p = 0.0016) (Figure 2D). The observed negative effects of resource evolution were driven by patterns in the second block, with weaker and somewhat opposite effects in block 1 (Figure S2).

**Table 4.**
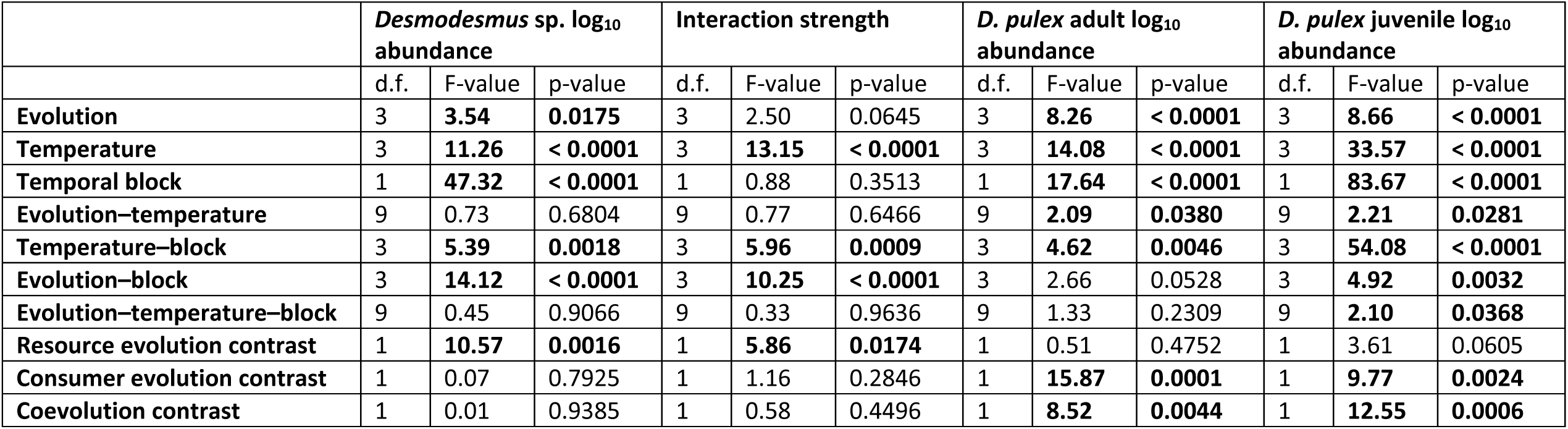
*Desmodesmus* sp. and *Daphnia pulex* univariate ANOVA (n = 8) results examining the effects of evolution of both the consumer and the resource in complex communities in response to warming. The model included an evolution treatment and temperature as two main effects, as well as a temporal blocking variable and all interactions. Here, the evolution treatment included the four possible combinations crossing the ambient- or warm-origin populations of both resources and consumers, but these analyses did not include algae-alone controls. The interaction between the evolution treatment and temperature evaluated any temperature-dependence of the effects of evolution. Orthogonal contrasts evaluated the effects of 1) resource evolution, 2) consumer evolution, and 3) coevolution between the resource and the consumer. Reported are Type III SS F-values and p-values for the model (due to one missing value causing a slightly unbalanced design). Significant results are in bold.

### Effects of evolutionary history on interaction strength

The importance of resource evolutionary history on algae abundance was reflected in the changes to interaction strength for each consumer–resource pair (Tables 3 and 4). For the *D. pulex*–*C. reinhardtii* pair, resource evolution reduced interaction strength (consumer–resource ANOVA resource evolution contrast: F_1_ = 54.90, p < 0.0001), but primarily at intermediate temperatures (interaction between evolutionary history and temperature: F_9_ = 5.16, p < 0.0001) (Figure 3A) in both blocks (Figure S2). For the *D. pulex*–*Desmodesmus* sp. pair, however, resource evolution increased interaction strength, independent of temperature (consumer–resource ANOVA resource evolution contrast: F_1_ = 5.86, p = 0.0174; interaction between evolutionary history and temperature: NS) (Figure 3B). For this species pair, overall trends in interaction strength were driven by patterns in block 2, here again with weaker and somewhat opposite effects in block 1 (Figure S2)

**Table 3.** *Chlamydomonas reinhardtii* and *Daphnia pulex* univariate ANOVA (n = 8) results examining the effects of evolution of both the consumer and the resource in complex communities in response to warming. The model included an evolution treatment and temperature as two main effects, as well as a temporal blocking variable and all interactions. Here, the evolution treatment included the four possible combinations crossing the ambient- or warm-origin populations of both resources and consumers, but these analyses did not include algae-alone controls. The interaction between the evolution treatment and temperature evaluated any temperature-dependence of the effects of evolution. Orthogonal contrasts evaluated the effects of 1) resource evolution, 2) consumer evolution, and 3) coevolution between the resource and the consumer. Reported are Type III SS F-values and p-values for the model (due to one missing value causing a slightly unbalanced design). Significant results are in bold.

|  | *C. reinhardtii* log<sub>10</sub><br>abundance |  |  | Interaction strength |  |  | *D. pulex* adult log<sub>10</sub><br>abundance |  |  | *D. pulex* juvenile log<sub>10</sub><br>abundance |  |  |
| --- | --- | --- | --- | --- | --- | --- | --- | --- | --- | --- | --- | --- |
|  | d.f. | F-value | p-value | d.f. | F-value | p-value | d.f. | F-value | p-value | d.f. | F-value | p-value |
| **Evolution** | 3 | **107.54** | **< 0.0001** | 3 | **19.64** | **< 0.0001** | 3 | 2.35 | 0.0772 | 3 | 1.80 | 0.1529 |
| **Temperature** | 3 | **17.48** | **< 0.0001** | 3 | **48.61** | **< 0.0001** | 3 | **6.37** | **0.0006** | 3 | **7.48** | **0.0001** |
| **Temporal block** | 1 | **162.18** | **< 0.0001** | 1 | **359.69** | **< 0.0001** | 1 | 0.10 | 0.7532 | 1 | 0.00 | 0.9543 |
| **Evolution–temperature** | 9 | **4.39** | **< 0.0001** | 9 | **5.16** | **< 0.0001** | 9 | 1.20 | 0.3067 | 9 | 0.53 | 0.8469 |
| **Temperature–block** | 3 | **12.52** | **< 0.0001** | 3 | **25.83** | **< 0.0001** | 3 | **5.08** | **0.0026** | 3 | 1.55 | 0.2062 |
| **Evolution–block** | 3 | **4.00** | **0.0099** | 3 | **11.85** | **< 0.0001** | 3 | **3.51** | **0.0183** | 3 | 2.31 | 0.0815 |
| **Evolution–temperature–block** | 9 | 1.63 | 0.1170 | 9 | **3.52** | **0.0009** | 9 | 1.40 | 0.2010 | 9 | 1.07 | 0.3949 |
| **Resource evolution contrast** | 1 | **318.47** | **< 0.0001** | 1 | **54.90** | **< 0.0001** | 1 | **5.50** | **0.0211** | 1 | **5.07** | **0.0266** |
| Consumer evolution contrast | 1 | 1.36 | 0.2456 | 1 | 1.36 | 0.2456 | 1 | 1.48 | 0.2262 | 1 | 0.29 | 0.5904 |
| Coevolution contrast | 1 | 2.53 | 0.1150 | 1 | 2.53 | 0.1150 | 1 | 0.01 | 0.9339 | 1 | 0.00 | 0.9740 |

**Figure 3A,B.**
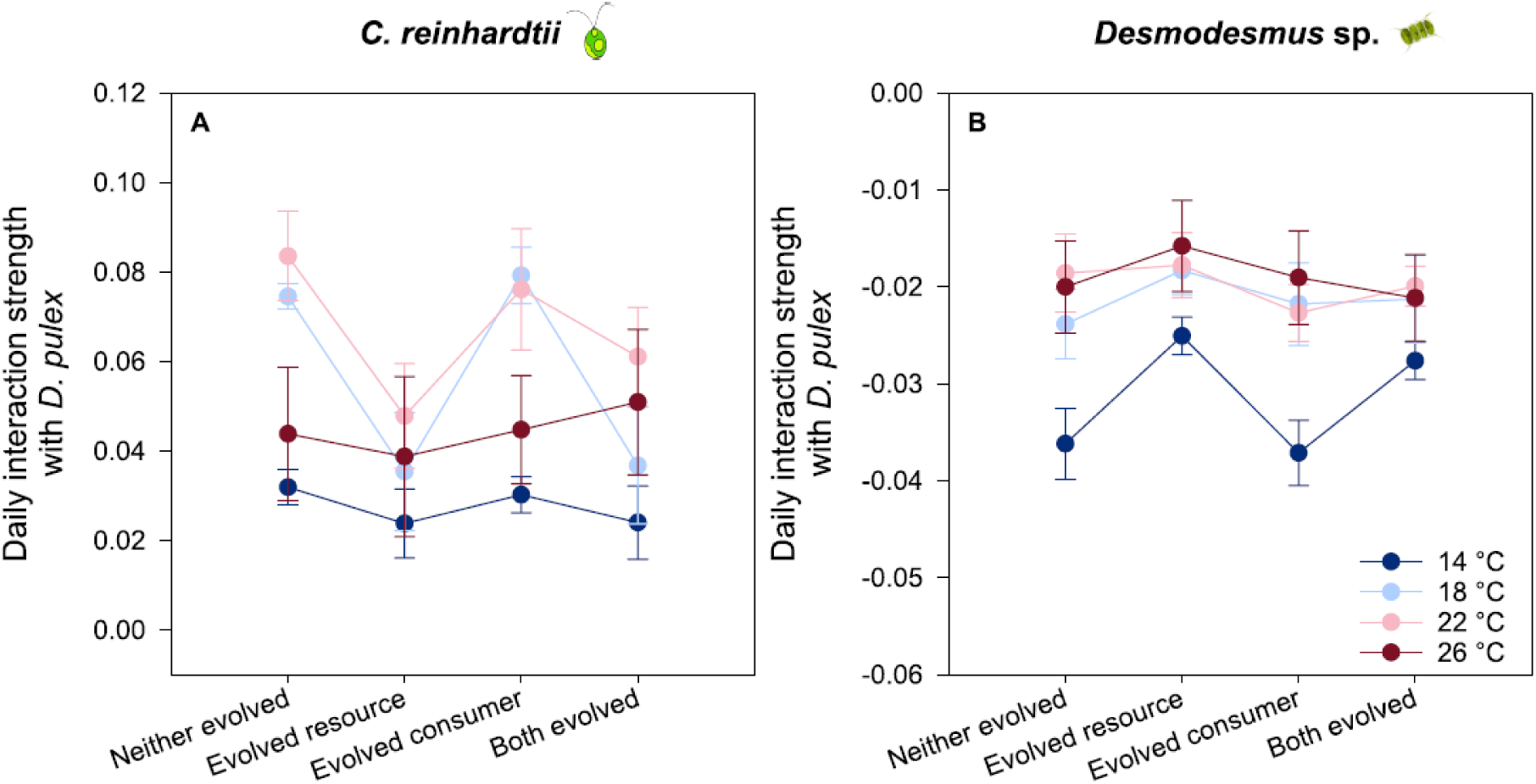
The effects of evolution in warmed communities on interaction strength of *C. reinhardtii* (A) and *Desmodesmus* sp. (B) with *D. pulex* depend upon current thermal. Evolutionary history treatments (where “evolved” refers to evolution in the warmed communities) are shown on the x-axis. Circles are the means (n = 8) of each treatment with the error bars showing ± the SEM. Colors correspond to temperature (dark blue = 14, light blue = 18, light pink = 22, and dark pink = 26 °C).

### Consumer responses to evolutionary history and temperature

For the consumer, *Daphnia pulex*, the effects of evolutionary history differed depending upon its resource species. When paired with *C. reinhardtii*, resource evolution significantly reduced the abundance of adult and juvenile *D. pulex* (adult resource evolution contrast: F_1_ = 5.50, p = 0.0211; juvenile resource evolution contrast: F_1_ = 5.07, p = 0.0266; Table 2, Figure 4A,B). The observed effect of resource evolution was primarily driven by patterns in block 1 (Figure S3). In this pairing, neither consumer evolution nor coevolution affected the abundance of *D. pulex* and the effects of resource evolution were independent of temperature (consumer–resource ANOVA interaction between evolutionary history and temperature: NS).

**Figure 4A–D.**
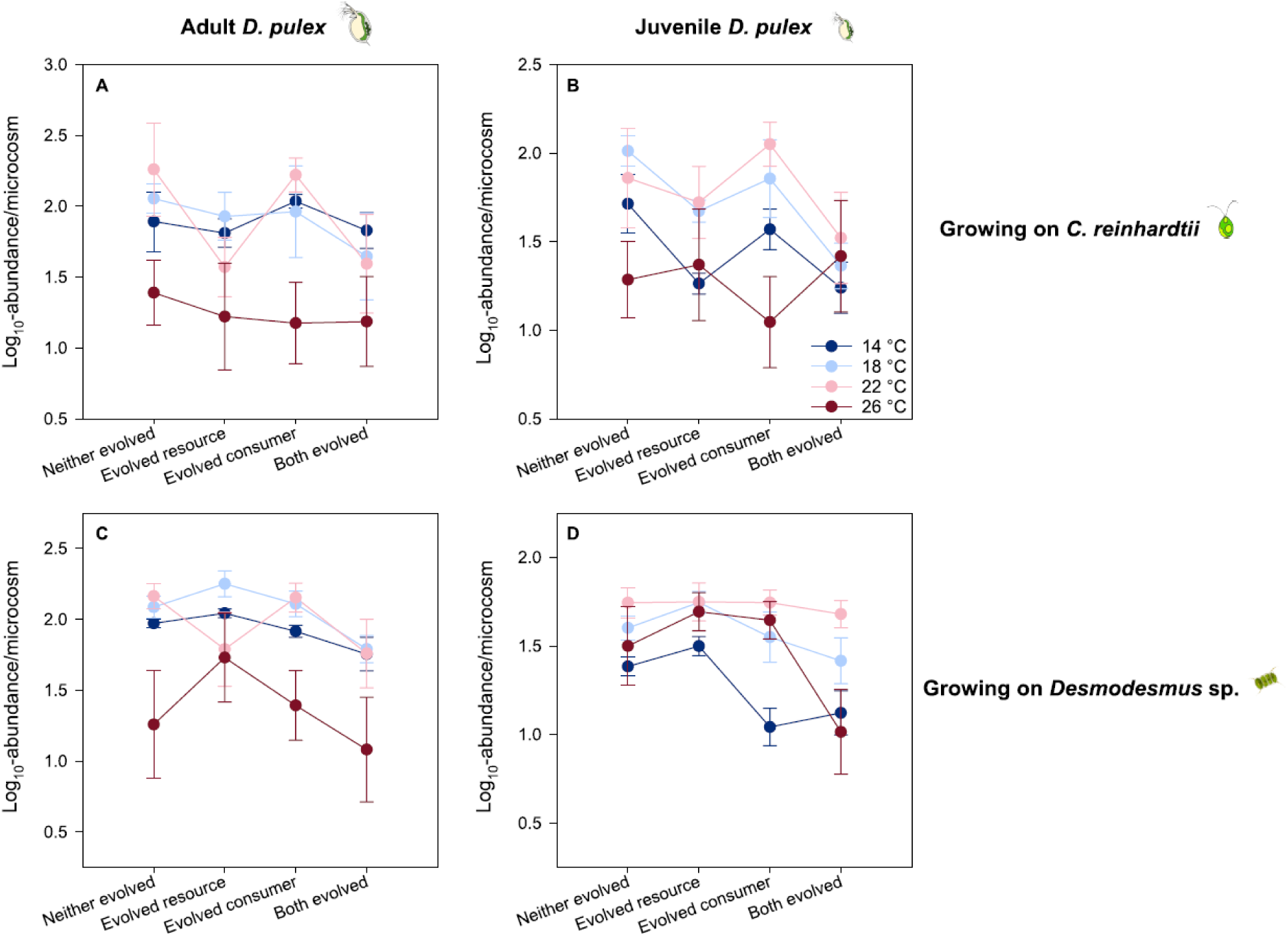
The effects of evolution in warmed communities on the log_10_-abundance of *D. pulex* juveniles (A) and adults (B) growing on *C. reinhardtii* and on *Desmodesmus* sp. (C, D) depend upon current thermal environment. In all plots, evolutionary history treatments (where “evolved” refers to evolution in the warmed communities) are shown on the x-axis. Circles are the means (n = 8) of each treatment with the error bars showing ± the SEM. Colors correspond to temperature (dark blue = 14, light blue = 18, light pink = 22, and dark pink = 26 °C.

When growing on the resource *Desmodesmus* sp., the effects of consumer evolution became apparent. In this case, warm-origin consumers had overall lower abundance than the ambient-origin population (adult consumer evolution contrast: F_1_ = 15.87, p = 0.0001; juvenile consumer evolution contrast: F_1_ = 9.77, p = 0.0024; Table 4, Figure 4C,D). There was no direct effect of *Desmodesmus* evolution, but coevolution between the resource and consumer affected consumer abundance (adult coevolution contrast: F_1_ = 8.52, p = 0.0044; juvenile coevolution contrast: F_1_ = 12.55, p = 0.0006). The effects on juveniles were driven in a large part by patterns in block 1 (Figure S3). The effects of evolutionary history were also dependent upon current thermal environment when growing on *Desmodesmus* sp. (evolutionary history-temperature interaction for adult and juvenile daphnids respectively: F_9_ = 2.09, p = 0.0380 and F_9_ = 2.21, p = 0.0281; Table 4, Figure 4C,D). For *D. pulex* growing on *Desmodesmus* sp., the effects of consumer evolution were most evident at 14 °C, where evolved *D. pulex* adults (“Evolved consumer” and “Both evolved” treatments in Figure 4) had significantly reduced abundance regardless of *Desmodesmus* evolutionary history compared to the ambient-origin *D. pulex* (“Neither evolved” and “Evolved resource” treatments). The effects of coevolution between the consumer and resource were strongest at 26 °C, with warm-adapted adult *D. pulex* abundance being significantly reduced only in the presence of warm-adapted *Desmodesmus* sp. (“Both evolved” treatment) (Figure 4D). In contrast to the temperature-dependence observed for the resource, *C. reinhardtii*, evolutionary history was less important for consumer abundance at intermediate temperatures.

### Robustness of the approach

Although we observed significant effects of temporal block, our planned contrasts examining the overall effects of evolutionary history lend us confidence in interpreting the overall importance of evolutionary history in driving our results. Additionally, because we both centered across temporal blocks, as well as examined patterns within blocks to understand interactions, we feel that our interpretation of our results is robust. The differences across temporal blocks could plausibly have resulted from a number of factors that we were unable to control completely. First, we attempted to maintain all algal source populations in the exponential phase (maximizing population growth rate at start) and standardize starting abundances. Nonetheless, starting populations differed somewhat in abundance within and across blocks. Similarly, some populations may have been in—or closer to— stationary phase than others, despite our efforts to maintain all at exponential growth. Similarly, although we standardized our selection of *Daphnia* by choosing the largest adults, adults may have differed in average age across the temporal blocks, potentially contributing to differences in *Daphnia* population size and growth rate. Finally, we swapped incubators among treatments across the two temporal blocks to avoid incubator pseudoreplication, possibly leading to unmeasured incubator effects.

## Discussion

Understanding the interaction between evolutionary history, the current abiotic environment, and biotic interactions is critical for a more nuanced understanding of how communities respond to anthropogenic stressors. Here we showed that both the current thermal environment, as well as the presence of interspecific interactions can affect the importance of prior evolution in response to warming in complex communities, and that this evolution also influences the outcomes of current ecological dynamics. For each consumer–resource pair, the effects of evolution were temperature-dependent, but both the effects and the temperature-dependence itself additionally depended upon the identity of evolving species in each pair.

### The importance of evolutionary history depends upon current thermal environment

Evolution in warmed communities had significant effects on the two algal resource species when grown alone, but these effects were opposite in direction and differed in magnitude. In one species, *Chlamydomonas reinhardtii*, prior evolution in warmed communities increased fitness (as assessed by abundance) across all temperatures, a result that is broadly consistent with the “hotter is better” hypothesis (Liu et al. 2022)(Figure S1), although our tested temperature range (determined by the thermal tolerance and activity of the consumer) precluded us from definitively evaluating competing proposed evolutionary responses. The positive direction of change agrees with a previous experiment examining the short-term effects of thermal evolution of *C. reinhardtii* independently isolated from these same mesocosms (Schaum et al. 2017). In the second alga, *Desmodesmus* sp., the effects of evolutionary history were dependent upon the current thermal environment. For this species, fitness was lower in the warm-origin genotype at three of the four tested temperatures, but the pattern was reversed at 14 °C, the lowest temperature in our range. This reversal of direction of effect across temperatures suggests the possibility of a response that is harmful in part of the thermal range, but possibly advantageous in another (Figure S1). The pattern of the result is surprising however, as the expectation is that warm-origin populations would perform better at warm temperatures compared to colder temperatures if the evolutionary response was directly driven by the experimentally imposed temperature treatment.

Evaluating the mechanisms underlying this result was impossible given that we only successfully isolated one genotype for each species from each of the ambient and warm mesocosms, however, these results may be consistent with one or more aspects of our system. First, our algal genotypes were isolated from the experimental ponds during the winter season, so this particular genotype may be more successful in the winter conditions experienced by the experimentally warmed ponds. Typical winter conditions in the ponds ranged from approximately 3–10 °C with a mean of around 6 °C (Yvon-Durocher et al. 2015), thus the additional +4 °C of warming experimentally imposed upon these ponds raised the winter temperature to within a few degrees of our 14 °C treatment. We did not examine the effects of evolutionary history at any temperature lower than 14 °C, however, so we are unable to compare the relative fitness of each genotype at typical winter temperatures compared to 14 °C. Conversely, the warm-origin genotype that we isolated may be maladaptive, with lower fitness than the ambient-origin genotype under most conditions. Anthropogenic warming is anticipated to create an extinction debt in many communities (Cotto et al. 2017), with poorly adapted species expected to be lost over time, especially as a result of stochastic processes combined with low dispersal (Kuussaari et al. 2009). It is possible that we captured part of this process, with a maladapted genotype “trapped” in the warm mesocosms. Another possibility is that the selection occurred in response to an agent that emerged secondary to the experimentally imposed temperature treatment. Our manipulative approach isolates temperature from other climatic variables that may affect evolutionary trajectory in natural populations, but our populations were nonetheless potentially evolving in response to a number of factors (e.g., community composition) that diverged over the course of this long-running experiment (Yvon-Durocher et al. 2011, 2015, Dossena et al. 2012). Thus, although we can ultimately link our responses to the experimentally-imposed warming treatment, populations are potentially evolving to a number of simultaneous and related stressors (terHorst et al. 2018). While this situation reflects the expectation for natural communities responding to habitat warming—namely that responses will encompass altered biotic interactions as well as direct responses to changed abiotic factors—it is therefore impossible for us to definitively disentangle the most likely mechanism driving the lowered fitness that we observed in warm-origin populations of *Desmodesmus* sp.

### Current ecological context modulates the temperature-dependence of the effects of evolutionary history

Our understanding of how evolution is altered in complex communities is still limited. However, previous studies suggest that evolution to environmental stressors, like habitat warming, that occurs in populations embedded within communities can yield different outcomes compared to evolution in single populations (Van Doorslaer et al. 2009b, 2010, terHorst et al. 2018, Hiltunen et al. 2018, De Meester et al. 2019). Our experiment complements a growing body of evidence that shows that evolutionary responses to warming do occur in complex communities (De Meester et al. 2019, McGaughran et al. 2021, Douce et al. 2024), and that such evolution has important consequences for both the direct responses to evolutionary stressors as well as for the interspecific interactions that ultimately shape communities.

The inclusion of consumer–resource interactions also introduced an additional layer of complexity necessary to understand how the importance of evolutionary history is affected by temperature. Organisms exhibit a mixture of unimodal and linear responses of traits to temperature, and interacting organisms differ in both the ecological and evolutionary responses of these traits. The interactions themselves may therefore depend upon temperature, influencing the outcomes of evolutionary change evaluated across temperatures (Barton and Yvon-Durocher 2019). For both consumer–resource pairs in our study system, the presence of the consumer altered the interaction between evolutionary history and temperature, indicating that examining the effects of past evolution in response to environmental change can be highly dependent on the current ecological conditions.

In the case of the *Daphnia pulex–C. reinhardtii* pair, the importance of evolutionary history was only temperature-dependent when the consumer was present. For this pair, the magnitude of the effect of evolutionary history on algal abundance was higher at intermediate temperatures compared to temperature extremes, adding nuance to the result that “hotter is better”, making it instead “hotter is best at moderate or intermediate temperatures”. Consumer evolution was unimportant when *D. pulex* was paired with *C. reinhardtii*—warm-origin *C. reinhardtii* significantly reduced the abundance of the consumer regardless of temperature. Thus, “hotter is better” for this resource translates into “hotter is worse” for the consumer in terms of the effects of resource evolution. We noted that the warm-origin genotype exhibited a mucilaginous extra-cellular matrix that might be expected to reduce ingestion rates for filter-feeding consumers like *D. pulex*. These results are also consistent with observations of an increased carbon to phosphorus ratio (i.e., reduced resource quality) observed previously in warm-adapted *C. reinhardtii* independently isolated from these experimental mesocosms (Yvon-Durocher et al. 2017). Such a reduction in resource quality would be expected to negatively affect consumer performance (Leal et al. 2017), but an analysis of nutrient content of the algae was outside the scope of our experiment. It is noteworthy, however, that despite the lack of a significant interaction between evolutionary history and temperature, the importance of resource evolution was strongest at 22 °C, the temperature at which *D. pulex* abundance was highest, and weakest at 26 °C, where the daphnids struggled to persist. The evolution that we observed in *C. reinhardtii* thus appeared to confer a double advantage to the resource, making it both more fit, while simultaneously reducing the success of the consumer, possibly the reason why we observed larger effect sizes for resource evolutionary history in the presence of the consumer, especially at intermediate temperatures (i.e., effect size more than doubled on a log_10_ scale).

The effects of consumer presence for *Desmodesmus* sp. were opposite to those for *C. reinhardtii*. Here, the presence of the consumer eliminated the temperature-dependence seen when *Desmodesmus* sp. grew alone, and even reversed the direction of the effect of evolutionary history at 14°C. When paired with the consumer, this subtlety across temperatures was lost, and warm-origin *Desmodesmus* sp. underperformed relative to ambient-origin *Desmodesmus* regardless of temperature. One possible explanation for the apparently reduced fitness of the warm-origin genotype could be that it served as a superior food source for the *Daphnia*. If this were the case, we might expect to see that the *Daphnia* achieved higher abundances when consuming the warm-origin algae. In fact, however, *Desmodesmus* sp. evolutionary history had contrasting effects on the consumer’s fitness depending upon consumer thermal origin (i.e., the effect of coevolution on *D. pulex* abundance in this pair).

Ambient-origin consumers exhibited a trend towards slightly higher fitness with warm-origin resource, but warm-origin *Daphnia* had lowest fitness when paired with warm-origin *Desmodesmus*. Thus, the effect does not appear to be consistent with increased consumption by the *Daphnia*. Additionally, in this pair, the *Daphnia* experienced temperature-dependent effects of evolutionary history. Here, effects were strongest at the two temperature extremes, with warm-origin *Daphnia* having lowest abundance at these temperatures, and especially at 26 °C and when paired with warm-origin *Desmodesmus* sp.

We examined the temperature-dependent consequences of evolution for two consumer– resource pairs over multiple generations of interactions and found evidence for adaptive evolution in response to warming in one of the two species pairs examined. For the *D. pulex–C. reinhardtii* pair, we observed adaptive evolution broadly consistent with expectations from the “hotter is better” hypothesis for the resource. While clearly, “hotter is better” did not apply to the consumer in our experiment, we did note a potential—albeit, non-significant—trend better matching “biochemical adaptation” for *D. pulex*, at least when paired with *C. reinhardtii*. Specifically, after examining just the performance of the warm-origin consumer relevant to the ambient-origin consumer when paired with the ambient-origin resource (i.e., removing resource evolution from the picture entirely), a trend emerged. The thermal optimum of *D. pulex* seemed to shift from 18 to 22 °C, without any increase in the magnitude of the optimum (Figure S2). Interestingly, however, no such trend is apparent when *D. pulex* is paired with *Desmodesmus* sp., nor does it seem to confer a measurable fitness advantage to the warm-origin consumer. Overall, hotter was worse from the perspective of the consumer, driven both by decreased performance in the presence of warm-origin resources (when paired with *C. reinhardtii*) and as a result of its own evolution (when paired with *Desmodesmus* sp.). These observations should be treated with caution, however, as the experimental design with four temperatures did not enable us to fit thermal performance curves to statistically evaluate these trends in the data. This result may seem surprising, but not only is evolutionary adaptation as a result of warming not expected in all species (Hoffmann and Sgró 2011), it also remains possible that the fitness consequences of the response depend upon the length of the temperature exposure. We focused on the effects of constant temperature over multiple generations, however, the semi-natural mesocosms from which we isolated these genotypes additionally fluctuate in temperature both diurnally and seasonally. These fluctuations may be an important component of the selection imposed by the experimental warming treatment (Diamond 2017), and organisms in warmed ponds may experience more frequent and intense, but shorter, periods of thermal stress. A more complete understanding of the interaction between current thermal environment and evolutionary history, therefore, will require an improved understanding of how populations respond to fluctuating temperatures.

## Conclusions

Our work demonstrates that local thermal environment and biotic interactions can together modulate the importance of past thermal evolutionary history. In some cases, evolution in response to warming can be beneficial for resources but detrimental for consumers, suggesting that we must account for the effects of warming on ingestion rates (see e.g., Barneche et al. 2021). In other cases, the picture is more complex and changes across a temperature range. This topic deserves further attention to improve our understanding of how populations and communities evolve in response to habitat warming, and how this then feeds back to drive the effects of the evolution. We have built upon previous studies that have primarily used space-for-time latitudinal substitutions in comparisons of different thermally adapted populations (e.g., Sanford et al. 2003, De Block et al. 2013, Dinh Van et al. 2013, Tran et al. 2016) and argue that future studies should consider both the current biotic and thermal context of their experiments, while definitely attributing evolutionary response to a manipulated factor of interest. These community properties can affect outcomes, not just in terms of the magnitude of the effect size, but even in direction of the effect. It is therefore of the utmost importance that we account for these interactions in our collective understanding. Evolution can drive responses in unexpected directions, but it is essential that we consider evolutionary responses to anthropogenic change in the context of both the current abiotic and biotic conditions to understand the full extent and direction of its effects.

## Acknowledgements

The French ANR through LabEx TULIP (ANR-10-LABX-41) and the FRAGCLIM Consolidator Grant, funded by the European Research Council under the European Union’s Horizon 2020 research and innovation programme (Grant Agreement Number 726176) supported this research.

## Author contributions

CAF and JMM conceived and designed the study. CAF, SAC, AG, and EB isolated organisms and collected data. CAF analyzed the data. CAF wrote the first draft of the manuscript. CAF, JMM, SAC, and EB jointly revised the manuscript. The authors declare no competing financial interests.

## References

1. Álvarez-Codesal, S., C. A. Faillace, A. Garreau, E. Bestion, A. D. Synodinos, and J. M. Montoya. 2023. Thermal mismatches explain consumer–resource dynamics in response to environmental warming. Ecology and Evolution 13:e10179.

2. Angilletta Jr., M. J., A. F. Bennett, H. Guderley, C. A. Navas, F. Seebacher, and R. S. Wilson. 2006. Coadaptation: A unifying principle in evolutionary thermal biology. Physiological and Biochemical Zoology 79:282–294.

3. Angilletta, M. J., R. B. Huey, and M. R. Frazier. 2010. Thermodynamic Effects on Organismal Performance: Is Hotter Better? Physiological and Biochemical Zoology 83:197–206.

4. Barneche, D. R., C. J. Hulatt, M. Dossena, D. Padfield, G. Woodward, M. Trimmer, and G. Yvon-Durocher. 2021. Warming impairs trophic transfer efficiency in a long-term field experiment. Nature 592:76–79.

5. Barton, S., and G. Yvon-Durocher. 2019. Quantifying the temperature dependence of growth rate in marine phytoplankton within and across species. Limnology and Oceanography 64:2081–2091.

6. Berlow, E. L., A.-M. Neutel, J. E. Cohen, P. C. De Ruiter, B. O. Ebenman, M. Emmerson, J. W. Fox, V. A. Jansen, J. I. Jones, and G. D. Kokkoris. 2004. Interaction strengths in food webs: issues and opportunities. Journal of animal ecology:585–598.

7. Comeau, A. M., G. M. Douglas, and M. G. I. Langille. 2017. Microbiome Helper: a Custom and Streamlined Workflow for Microbiome Research. mSystems.

8. Cotto, O., J. Wessely, D. Georges, G. Klonner, M. Schmid, S. Dullinger, W. Thuiller, and F. Guillaume. 2017. A dynamic eco-evolutionary model predicts slow response of alpine plants to climate warming. Nature Communications 8:15399.

9. Craddock, D. R. 1976. Effects of increased water temperature on *Daphnia pulex*. Fishery Bulletin 74:403– 408.

10. De Block, M., K. Pauwels, M. Van Den Broeck, L. De Meester, and R. Stoks. 2013. Local genetic adaptation generates latitude-specific effects of warming on predator-prey interactions. Global Change Biology 19:689–696.

11. De Meester, L., K. I. Brans, L. Govaert, C. Souffreau, S. Mukherjee, H. Vanvelk, K. Korzeniowski, L. Kilsdonk, E. Decaestecker, R. Stoks, and M. C. Urban. 2019. Analysing eco-evolutionary dynamics—The challenging complexity of the real world. Functional Ecology 33:43–59.

12. Dell, A. I., S. Pawar, and V. M. Savage. 2011. Systematic variation in the temperature dependence of physiological and ecological traits. Proceedings of the National Academy of Sciences 108:10591– 10596.

13. DeLong, J. P., G. Bachman, J. P. Gibert, T. M. Luhring, K. L. Montooth, A. Neyer, and B. Reed. 2018. Habitat, latitude and body mass influence the temperature dependence of metabolic rate. Biology Letters 14:20180442.

14. Diamond, S. E. 2017. Evolutionary potential of upper thermal tolerance: biogeographic patterns and expectations under climate change. Annals of the New York Academy of Sciences 1389:5–19.

15. Diamond, S. E., L. Chick, A. Perez, S. A. Strickler, and R. A. Martin. 2017. Rapid evolution of ant thermal tolerance across an urban-rural temperature cline. Biological Journal of the Linnean Society 121:248–257.

16. Dinh Van, K., L. Janssens, S. Debecker, M. De Jonge, P. Lambret, V. Nilsson-Örtman, L. Bervoets, and R. Stoks. 2013. Susceptibility to a metal under global warming is shaped by thermal adaptation along a latitudinal gradient. Global Change Biology 19:2625–2633.

17. Dossena, M., G. Yvon-Durocher, J. Grey, J. M. Montoya, D. M. Perkins, M. Trimmer, and G. Woodward. 2012. Warming alters community size structure and ecosystem functioning. Proceedings of the Royal Society B: Biological Sciences 279:3011–3019.

18. Douce, P., L. Simon, F. Colas, F. Mermillod-Blondin, D. Renault, C. Sulmon, P. Eymar-Dauphin, R. Dubreucque, and A.-K. Bittebiere. 2024. Warming drives feedback between plant phenotypes and ecosystem functioning in sub-Antarctic ponds. Science of The Total Environment 914:169504.

19. Faillace, C. A., A. Sentis, and J. M. Montoya. 2021. Eco-evolutionary consequences of habitat warming and fragmentation in communities. Biological Reviews 96:1933–1950.

20. Gaitán-Espitia, J. D., and A. J. Hobday. 2021. Evolutionary principles and genetic considerations for guiding conservation interventions under climate change. Global Change Biology 27:475–488.

21. Geerts, A. N., J. Vanoverbeke, B. Vanschoenwinkel, W. Van Doorslaer, H. Feuchtmayr, D. Atkinson, B. Moss, T. A. Davidson, C. D. Sayer, and L. De Meester. 2015. Rapid evolution of thermal tolerance in the water flea *Daphnia*. Nature Climate Change 5:665.

22. Gilchrist, G. W., and R. B. Huey. 1999. The direct response of *Drosophila melanogaster* to selection on knockdown temperature. Heredity 83:15–29.

23. Hague, M. T. J., and E. J. Routman. 2016. Does population size affect genetic diversity? A test with sympatric lizard species. Heredity 116:92–98.

24. Hangartner, S., and A. A. Hoffmann. 2016. Evolutionary potential of multiple measures of upper thermal tolerance in *Drosophila melanogaster*. Functional Ecology 30:442–452.

25. Higgins, J. K., H. J. MacLean, L. B. Buckley, and J. G. Kingsolver. 2014. Geographic differences and microevolutionary changes in thermal sensitivity of butterfly larvae in response to climate. Functional Ecology 28:982–989.

26. Hiltunen, T., J. Cairns, J. Frickel, M. Jalasvuori, J. Laakso, V. Kaitala, S. Künzel, E. Karakoc, and L. Becks. 2018. Dual-stressor selection alters eco-evolutionary dynamics in experimental communities. Nature Ecology and Evolution 2:1974–1981.

27. Hoffmann, A. A., and C. M. Sgró. 2011. Climate change and evolutionary adaptation. Nature 470:479– 485.

28. Huey, R. B., and J. G. Kingsolver. 2011. Variation in universal temperature dependence of biological rates. Proceedings of the National Academy of Sciences 108:10377–10378.

29. Johansson, M. P., F. Ermold, B. K. Kristjánsson, and A. Laurila. 2016. Divergence of gastropod life history in contrasting thermal environments in a geothermal lake. Journal of Evolutionary Biology 29:2043–2053.

30. Johansson, M. P., and A. Laurila. 2017. Maximum thermal tolerance trades off with chronic tolerance of high temperature in contrasting thermal populations of *Radix balthica*. Ecology and Evolution 7:3149–3156.

31. Kingsolver, J. G., and R. B. Huey. 2008. Size, temperature, and fitness: three rules. Evolutionary Ecology Research 10:251–268.

32. Knies, J. L., R. Izem, K. L. Supler, J. G. Kingsolver, and C. L. Burch. 2006. The genetic basis of thermal reaction norm evolution in lab and natural phage populations. PLoS Biology 4:1257–1264.

33. Kuussaari, M., R. Bommarco, R. K. Heikkinen, A. Helm, J. Krauss, R. Lindborg, E. Öckinger, M. Pärtel, J. Pino, F. Rodà, C. Stefanescu, T. Teder, M. Zobel, and I. Steffan-Dewenter. 2009. Extinction debt: a challenge for biodiversity conservation. Trends in Ecology & Evolution 24:564–571.

34. Latimer, C. A. L., R. S. Wilson, and S. F. Chenoweth. 2011. Quantitative genetic variation for thermal performance curves within and among natural populations of Drosophila serrata. Journal of Evolutionary Biology 24:965–975.

35. Leal, M. C., O. Seehausen, and B. Matthews. 2017. The Ecology and Evolution of Stoichiometric Phenotypes. Trends in Ecology & Evolution 32:108–117.

36. Liu, K., B. Chen, and H. Liu. 2022. Evidence of partial thermal compensation in natural phytoplankton assemblages. Limnology and Oceanography Letters 7:122–130.

37. Logan, M. L., R. M. Cox, and R. Calsbeek. 2014. Natural selection on thermal performance in a novel thermal environment. Proceedings of the National Academy of Sciences of the United States of America 111:14165–9.

38. Lurgi, M., B. C. López, and J. M. Montoya. 2012. Novel communities from climate change. Philosophical Transactions of the Royal Society B: Biological Sciences 367:2913–2922.

39. McGaughran, A., R. Laver, and C. Fraser. 2021. Evolutionary Responses to Warming. Trends in Ecology & Evolution 36:591–600.

40. O’Connor, M. I., M. F. Piehler, D. M. Leech, A. Anton, and J. F. Bruno. 2009. Warming and resource availability shift food web structure and metabolism. PLoS biology 7:e1000178.

41. O’Connor, M. I., E. R. Selig, M. L. Pinsky, and F. Altermatt. 2012. Toward a conceptual synthesis for climate change responses. Global Ecology and Biogeography 21:693–703.

42. O’Donnell, D. R., C. R. Hamman, E. C. Johnson, C. T. Kremer, C. A. Klausmeier, and E. Litchman. 2018. Rapid thermal adaptation in a marine diatom reveals constraints and trade-offs. Global Change Biology 24:4554–4565.

43. Padfield, D., G. Yvon-Durocher, A. Buckling, S. Jennings, and G. Yvon-Durocher. 2016. Rapid evolution of metabolic traits explains thermal adaptation in phytoplankton. Ecology Letters 19:133–142.

44. Parmesan, C. 2006. Ecological and Evolutionary Responses to Recent Climate Change. Annual Review of Ecology, Evolution, and Systematics 37:637–669.

45. Radchuk, V., T. Reed, C. Teplitsky, M. van de Pol, A. Charmantier, C. Hassall, P. Adamík, F. Adriaensen, M. P. Ahola, P. Arcese, J. Miguel Avilés, J. Balbontin, K. S. Berg, A. Borras, S. Burthe, J. Clobert, N. Dehnhard, F. de Lope, A. A. Dhondt, N. J. Dingemanse, H. Doi, T. Eeva, J. Fickel, I. Filella, F. Fossøy, A. E. Goodenough, S. J. G. Hall, B. Hansson, M. Harris, D. Hasselquist, T. Hickler, J. Joshi, H. Kharouba, J. G. Martínez, J.-B. Mihoub, J. A. Mills, M. Molina-Morales, A. Moksnes, A. Ozgul, D. Parejo, P. Pilard, M. Poisbleau, F. Rousset, M.-O. Rödel, D. Scott, J. C. Senar, C. Stefanescu, B. G. Stokke, T. Kusano, M. Tarka, C. E. Tarwater, K. Thonicke, J. Thorley, A. Wilting, P. Tryjanowski, J. Merilä, B. C. Sheldon, A. Pape Møller, E. Matthysen, F. Janzen, F. S. Dobson, M. E. Visser, S. R. Beissinger, A. Courtiol, and S. Kramer-Schadt. 2019. Adaptive responses of animals to climate change are most likely insufficient. Nature Communications 10:3109.

46. Rezende, E. L., L. E. Castañeda, and M. Santos. 2014. Tolerance landscapes in thermal ecology. Functional Ecology 28:799–809.

47. Rudman, S. M., M. Kreitzman, K. M. A. Chan, and D. Schluter. 2017. Evosystem Services: Rapid Evolution and the Provision of Ecosystem Services. Trends in Ecology & Evolution 32:403–415.

48. Sanford, E., M. S. Roth, G. C. Johns, J. P. Wares, and G. N. Somero. 2003. Local selection and latitudinal variation in a marine predator-prey interaction. Science 300:1135–1137.

49. SAS Institute. 2011. The SAS system for Windows. SAS Institute, Cary, NC.

50. Schaum, C.-E., S. Barton, E. Bestion, A. Buckling, B. Garcia-Carreras, P. Lopez, C. Lowe, S. Pawar, N. Smirnoff, M. Trimmer, and G. Yvon-Durocher. 2017. Adaptation of phytoplankton to a decade of experimental warming linked to increased photosynthesis. Nature Ecology and Evolution 1:94.

51. Schielzeth, H. 2010. Simple means to improve the interpretability of regression coefficients. Methods Ecol Evol 1: 103–113.

52. Tabi, A., F. Pennekamp, F. Altermatt, R. Alther, E. A. Fronhofer, K. Horgan, E. Mächler, M. Pontarp, O. L. Petchey, and S. Saavedra. 2020. Species multidimensional effects explain idiosyncratic responses of communities to environmental change. Nature Ecology and Evolution:1–8.

53. Terhorst, C. P., T. E. Miller, and D. R. Levitan. 2010. Evolution of prey in ecological time reduces the effect size of predators in experimental microcosms. Ecology 91:629–636.

54. terHorst, C. P., P. C. Zee, K. D. Heath, T. E. Miller, A. I. Pastore, S. Patel, S. J. Schreiber, M. J. Wade, and M. R. Walsh. 2018. Evolution in a community context: Trait responses to multiple species interactions. The American Naturalist 191:368–380.

55. Tran, T. T., L. Janssens, K. V. Dinh, L. Op de Beeck, and R. Stoks. 2016. Evolution determines how global warming and pesticide exposure will shape predator–prey interactions with vector mosquitoes. Evolutionary Applications 9:818–830.

56. Van Doorslaer, W., R. Stoks, C. Duvivier, A. Bednarska, and L. De Meester. 2009a. Population dynamics determine genetic adaptation to temperature in *Daphnia*. Evolution 63:1867–1878.

57. Van Doorslaer, W., R. Stoks, I. Swillen, H. Feuchtmayr, D. Atkinson, B. Moss, and L. De Meester. 2010. Experimental thermal microevolution in community-embedded *Daphnia* populations. Climate Research 43:81–89.

58. Van Doorslaer, W., J. Vanoverbeke, C. Duvivier, S. Rousseaux, M. Jansen, B. Jansen, H. Feuchtmayr, D. Atkinson, B. Moss, R. Stoks, and L. De Meester. 2009b. Local adaptation to higher temperatures reduces immigration success of genotypes from a warmer region in the water flea *Daphnia*. Global Change Biology 15:3046–3055.

59. Yvon-Durocher, G., A. P. Allen, M. Cellamare, M. Dossena, K. J. Gaston, M. Leitao, J. M. Montoya, D. C. Reuman, G. Woodward, and M. Trimmer. 2015. Five years of experimental warming increases the biodiversity and productivity of phytoplankton. PLoS Biology 13:e1002324.

60. Yvon-Durocher, G., J. M. Montoya, M. Trimmer, and G. Woodward. 2011. Warming alters the size spectrum and shifts the distribution of biomass in freshwater ecosystems. Global Change Biology 17:1681–1694.

61. Yvon-Durocher, G., C.-E. Schaum, and M. Trimmer. 2017. The temperature dependence of phytoplankton stoichiometry: investigating the roles of species sorting and local adaptation. Frontiers in Microbiology 8:2003.

