## Supplemental Figures S1-S4 for "Thermal environment and ecological interactions modulate the importance of evolution in response to warming"

**Supporting information for "Thermal environment and ecological interactions modulate the importance of evolution in response to warming"**

Cara A. Faillace, Soraya Álvarez-Codesal, Alexandre Garreau, Elvire Bestion, and José M. Montoya

**Supplemental Figure S1**. Patterns of algae abundance across temperatures depended upon the evolutionary history in response to prior warming ("evolved") of both resources and consumers for both *C. reinhardtii* (A) and *Desmodesmus* sp. (B). Circles are the means (n = 8) of each treatment with the error bars showing ± the SEM.

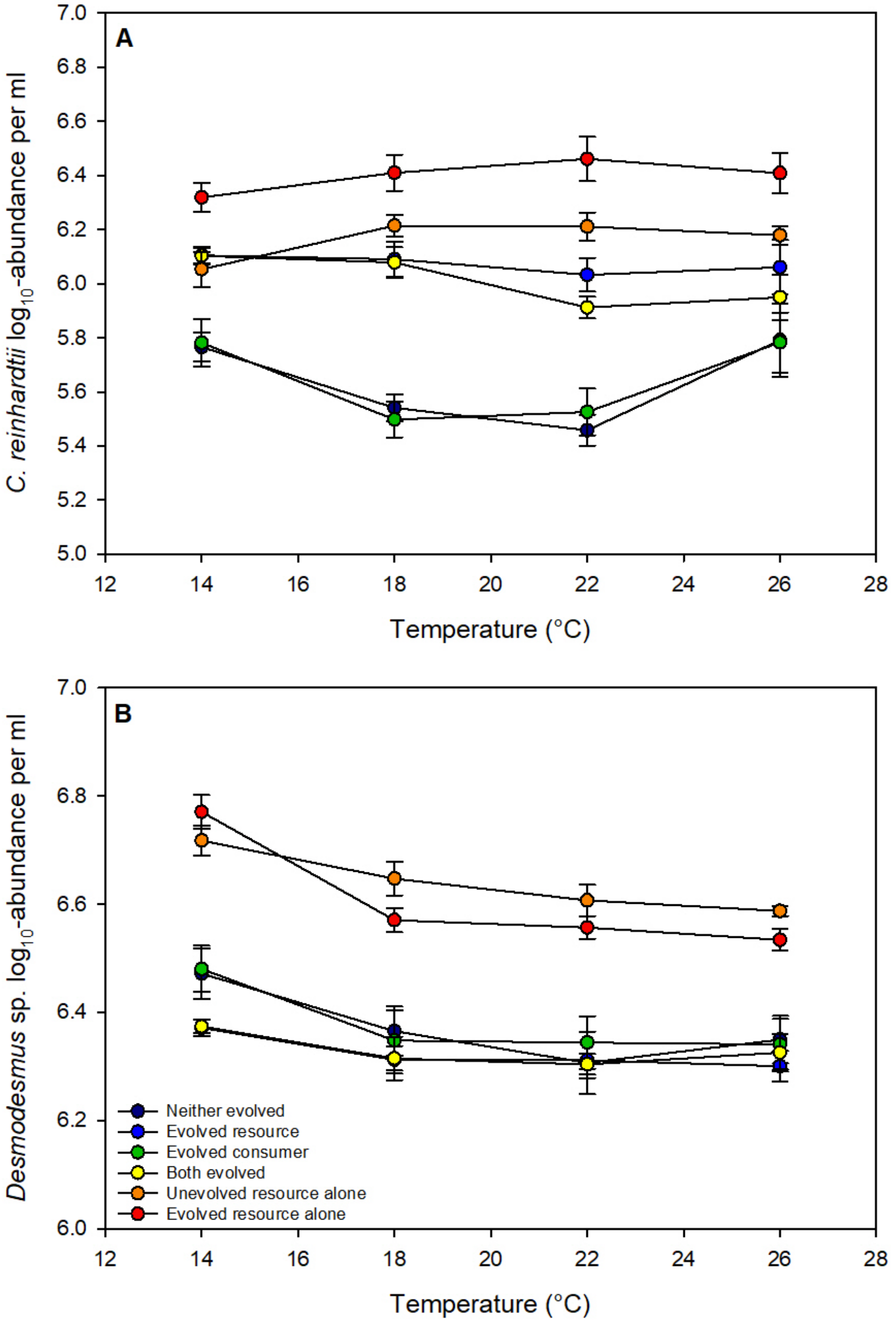

**Supporting information for "Thermal environment and ecological interactions modulate the importance of evolution in response to warming"**

Cara A. Faillace, Soraya Álvarez-Codesal, Alexandre Garreau, Elvire Bestion, and José M. Montoya

**Supplemental Figure S2**. Block effects and block interactions occurred for both species of algae when grown in the absence of consumers (A,B) and in presence of consumers (C–E). See Tables 1–4 (main text) for significance of block effects and block interactions. Evolutionary history treatments ("evolved" refers to evolution in the warmed communities) are on the x-axis. Circles are treatment means (n = 8) with the error bars showing ± the SEM. Colors correspond to temperature (dark blue = 14, light blue = 18, light pink = 22, and dark pink = 26 °C). Solid lines are for block 1 and dashed lines for block 2.

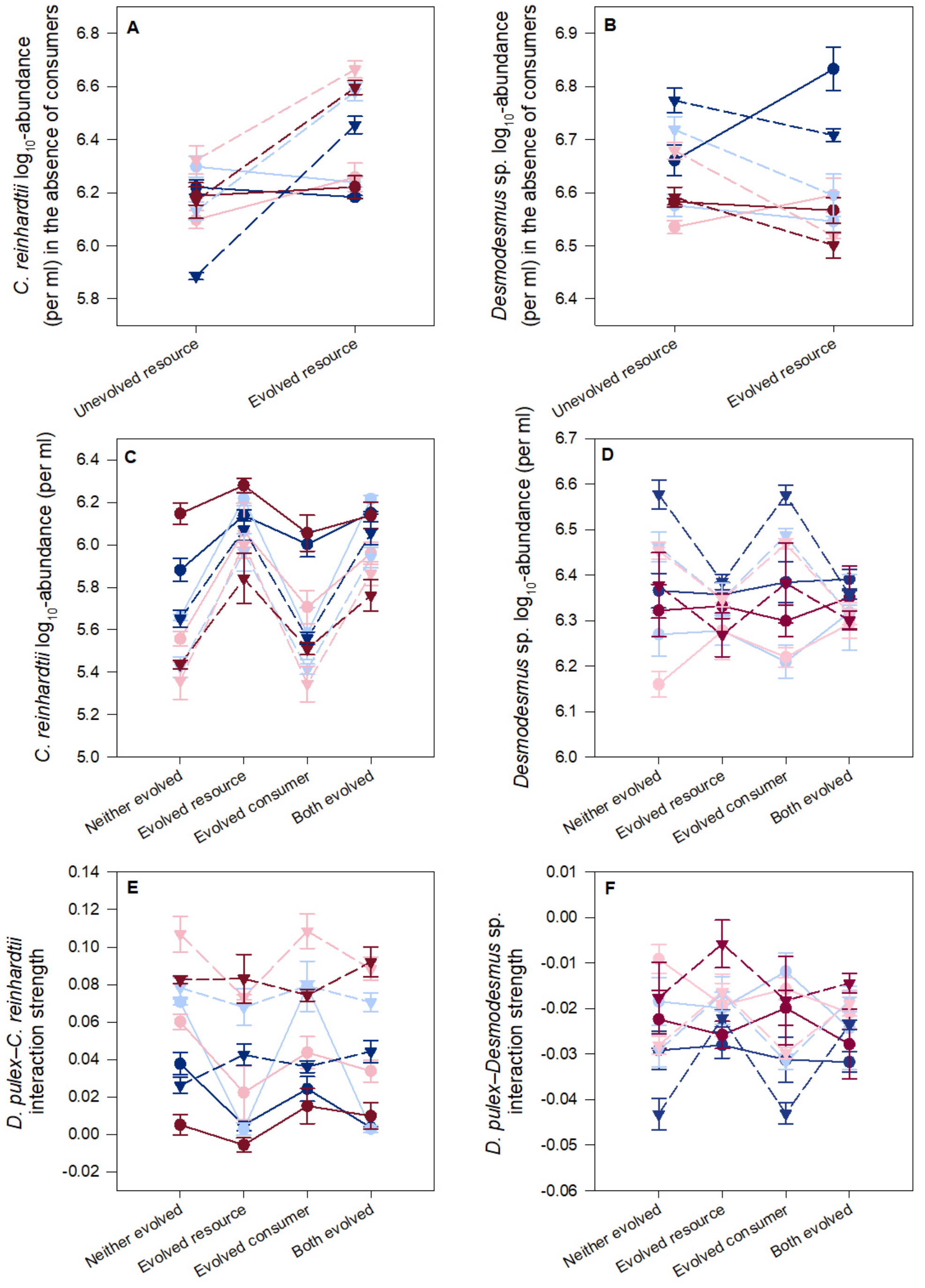

**Supporting information for "Thermal environment and ecological interactions modulate the importance of evolution in response to warming"**

Cara A. Faillace, Soraya Álvarez-Codesal, Alexandre Garreau, Elvire Bestion, and José M. Montoya

**Supplemental Figure S3**. Block effects and block interactions occurred for *D. pulex* when grown on *C. reinhardtii* (A, B) or *Desmodesmus* sp. (C, D). See the main text for significance of block effects and block interactions. Evolutionary history treatments ("evolved" refers to evolution in the warmed communities) are shown on the x-axis. Circles are the treatment means (n = 8) with the error bars showing ± the SEM. Colors correspond to temperature (dark blue = 14, light blue = 18, light pink = 22, and dark pink = 26 °C). Solid lines are for block 1 and dashed lines for block 2.

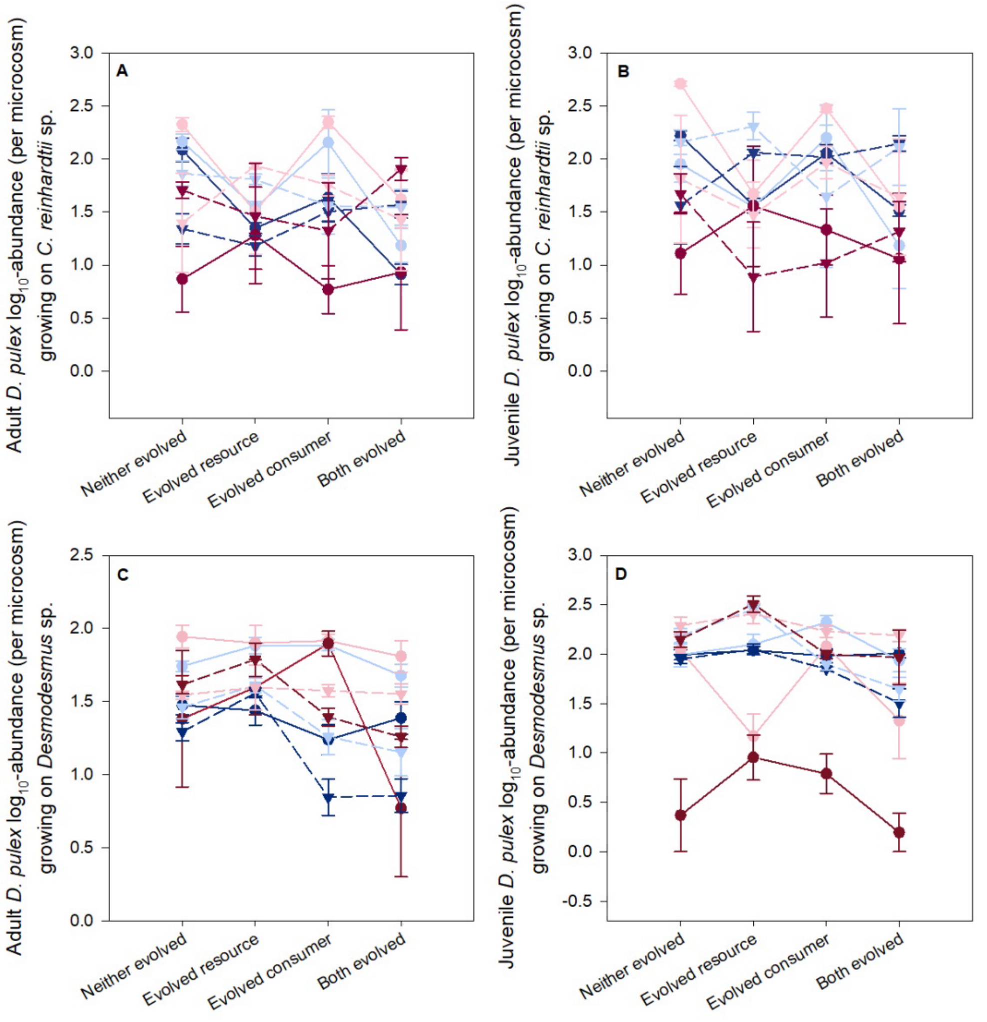

**Supporting information for "Thermal environment and ecological interactions modulate the importance of evolution in response to warming"**

Cara A. Faillace, Soraya Álvarez-Codesal, Alexandre Garreau, Elvire Bestion, and José M. Montoya

**Supplemental Figure S4**. Patterns of abundance of *D. pulex* across temperatures when grown on *C. reinhardtii* (A, B) or *Desmodesmus* sp. (D,C) depended upon the evolutionary history in response to prior warming ("evolved"). Circles are the treatment means (n = 8) with the error bars showing ± the SEM.

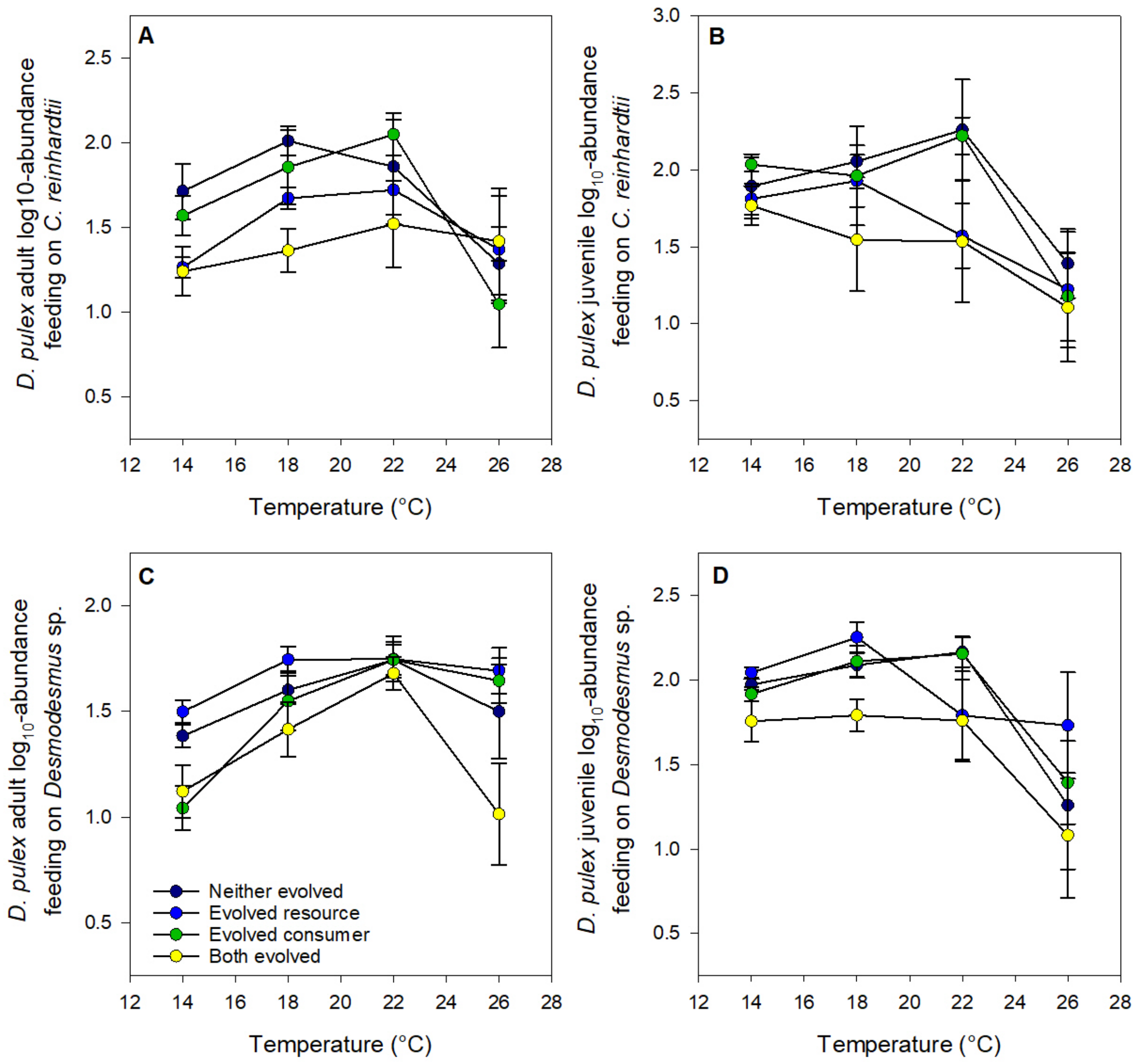
